# CWL-Airflow: a lightweight pipeline manager supporting Common Workflow Language

**DOI:** 10.1101/249243

**Authors:** Michael Kotliar, Andrey V. Kartashov, Artem Barski

## Abstract

**Background:** Massive growth in the amount of research data and computational analysis has led to increased utilization of pipeline managers in biomedical computational research. However, each of more than 100 such managers uses its own way to describe pipelines, leading to difficulty porting workflows to different environments and therefore poor reproducibility of computational studies. For this reason, the Common Workflow Language (CWL) was recently introduced as a specification for platform-independent workflow description, and work began to transition existing pipelines and workflow managers to CWL.

**Findings:** Here, we present CWL-Airflow, an extension for the Apache Airflow pipeline manager supporting CWL. CWL-Airflow utilizes CWL v1.0 specification and can be used to run workflows on standalone MacOS/Linux servers, on clusters, or on a variety of cloud platforms. A sample CWL pipeline for processing of ChIP-Seq data is provided.

**Conclusions:** CWL-Airflow will provide users with the features of a fully-fledged pipeline manager and an ability to execute CWL workflows anywhere Airflow can run—from a laptop to cluster or cloud environment.

**Availability:** CWL-Airflow is available under Apache license v.2 and can be downloaded from https://barski-lab.github.io/cwl-airflow, http://doi.org/10.5281/zenodo.2669582, RRID: SCR_017196.

## Background

Modern biomedical research has seen a remarkable increase in the production and computational analysis of large datasets, leading to an urgent need to share standardized analytical techniques. However, of the more than one hundred computational workflow systems used in biomedical research, most define their own specifications for computational pipelines [1,2]. Furthermore, the evolving complexity of computational tools and pipelines makes it nearly impossible to reproduce computationally heavy studies or to repurpose published analytical workflows. Even when the tools are published, the lack of a precise description of the operating system environment and component software versions can lead to inaccurate reproduction of the analyses—or analyses failing altogether when executed in a different environment. To ameliorate this situation, a team of researchers and software developers formed the Common Workflow Language (CWL) working group [3] with the intent of establishing a specification for describing analysis workflows and tools in a way that makes them portable and scalable across a variety of software and hardware environments. The CWL specification provides a set of formalized rules that can be used to describe each command line tool and its parameters, and optionally a container (e.g., a Docker [4] or Singularity [5] image) with the tool already installed. CWL workflows are composed of one or more of such command line tools. Thus, CWL provides a description of the working environment and version of each tool, how the tools are “connected” together, and what parameters were used in the pipeline. Researchers using CWL are then able to deposit descriptions of their tools and workflows into a repository (e.g., dockstore.org) upon publication, thus making their analyses reusable by others.

After version 1.0 of the CWL standard [6] and the reference executor, cwl-tool, were finalized in 2016, developers began adapting the existing pipeline managers to use CWL. For example, companies such as Seven Bridges Genomics and Curoverse are developing the commercial platforms Rabix [7] and Arvados [8] whereas academic developers (*e.g.*, Galaxy [9], Toil [10] and others) are adding CWL support to their pipeline managers (See Table 1 for the comparison of their features).

**Table 1.** CWL-Airflow and cwltool average execution time (seconds ± SEM^1^, n=3).

| Pipeline | CWL-Airflow |  | Cwltool |
| --- | --- | --- | --- |
|  | 1 node<br>1 workflow at a time | 3 nodes<br>3 workflows at a time | 1 node<br>1 workflow at a time |
| BioWardrobe ChIP-Seq Workflow | 1141 ± 18 | 1231 ± 3 | 955 ± 1 |
| ENCODE ChIP-Seq Mapping Workflow | 3784 ± 10 | 3824 ± 28 | 3245 ± 7 |

<sup>1</sup> standard error of the mean

Airflow [11] is a lightweight workflow manager initially developed by AirBnB, which is currently an Apache Incubator project, and is available under a permissive Apache license. Airflow executes each workflow as a Directed Acyclic Graph (DAG) of tasks that have directional noncircular dependencies. Tasks are usually atomic and are not supposed to share any resources with each other; therefore, they can be run independently. DAG describes relationships between the tasks and defines their execution order. DAG objects are initiated from Python scripts placed in a designated folder. Airflow has a modular architecture and can distribute tasks to an arbitrary number of workers, across multiple servers, while adhering to the task sequence and dependencies specified in the DAG. Unlike many of the more complicated platforms, Airflow imposes little overhead, is easy to install, and can be used to run task-based workflows in various environments ranging from standalone desktops and servers to Amazon or Google cloud platforms. It also scales horizontally on clusters managed by Apache Mesos [12] and may be configured to send tasks to Celery [13] task queue. Here we present an extension of Airflow, allowing it to run CWL-based pipelines. Altogether, this gives us a lightweight workflow management system with full support for CWL, the most promising scientific workflow description language.

## Methods

The CWL-Airflow package extends Airflow’s functionality with the ability to parse and execute workflows written with the current CWL v1.0 specification [6]. CWL-Airflow can be easily integrated into the Airflow scheduler logic as shown in the structure diagram in Figure 1. The Apache Airflow code is extended with a Python package that defines four basic classes—*CWLStepOperator, JobDispatcher, JobCleanup*, and *CWLDAG.* Additionally, the automatically generated *cwl_dag.py* script is placed in the DAGs folder. While periodically loading DAGs from the DAGs folder the Airflow scheduler runs the *cwl_dag.py* script and creates DAGs based on the available jobs and corresponding CWL workflow descriptor files.

**Figure 1.**
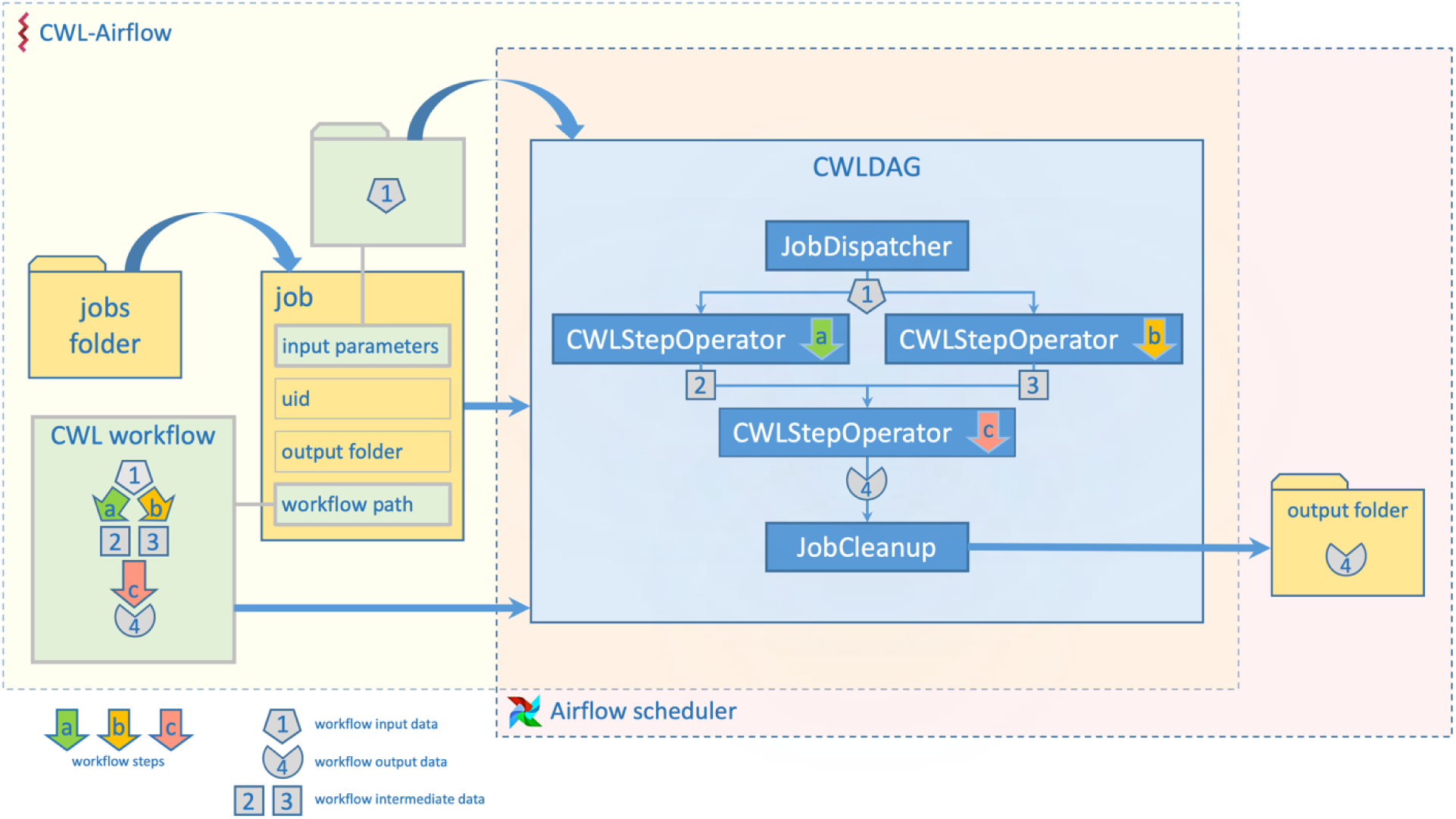
CWL-Airflow diagram. Job file contains information about CWL workflow and inputs. CWL-Airflow creates CWLDAG class instance based on the workflow structure and executes it in Airflow. The results are saved to the output folder.

In order to run a CWL workflow in Airflow, a file describing the job should be placed in the *jobs* folder (Fig. 1). Jobs are described by a text file (JSON or YAML) that includes workflow specific *input parameters* (e.g. input file locations) and three mandatory fields: *workflow* (absolute path to the CWL descriptor file to be run with this job), *output_folder* (absolute path to the folder where all the output files should be moved after successful pipeline execution) and *uid* (unique identifier for the run). CWL-Airflow parses every job file from the *jobs* folder, loads corresponding CWL workflow descriptor file and creates CWLDAG class instance based on the workflow structure and *input parameters* provided in the job file. The *uid* field from the job file is used to identify the newly created CWLDAG class instance.

CWLDAG is a class for combining the tasks into the DAG that reflects the CWL workflow structure. Every *CWLStepOperator* task corresponds to the workflow step and depends on others based on the workflow step inputs and outputs. This implements dataflow principles and architecture that are missing in Airflow. Additionally, *JobDispatcher* and *JobCleanup* tasks are added to the graph. *JobDisptacher* is used to serialize the *input parameters* from the job file and provide the pipeline with the input data; *JobCleanup* returns the calculated results to the *output folder*. When the Airflow scheduler executes the pipeline from the CWLDAG it runs the workflow with the structure identical to the CWL descriptor file used to create this graph.

While running CWL-Airflow on a single node might be sufficient in most of the cases, when it comes to the computationally intensive pipelines it is worth switching to the multi-node configuration (Fig. 2). Airflow uses Celery task queue to distribute processing over the multiple nodes. Celery provides the mechanisms for queueing and assigning tasks to the multiple workers, whereas the Airflow scheduler uses Celery executor to submit tasks to the queue. The Celery system helps to not only balance the load over the different machines, but also to define task priorities by assigning them to the separate queues.

**Figure 2.**
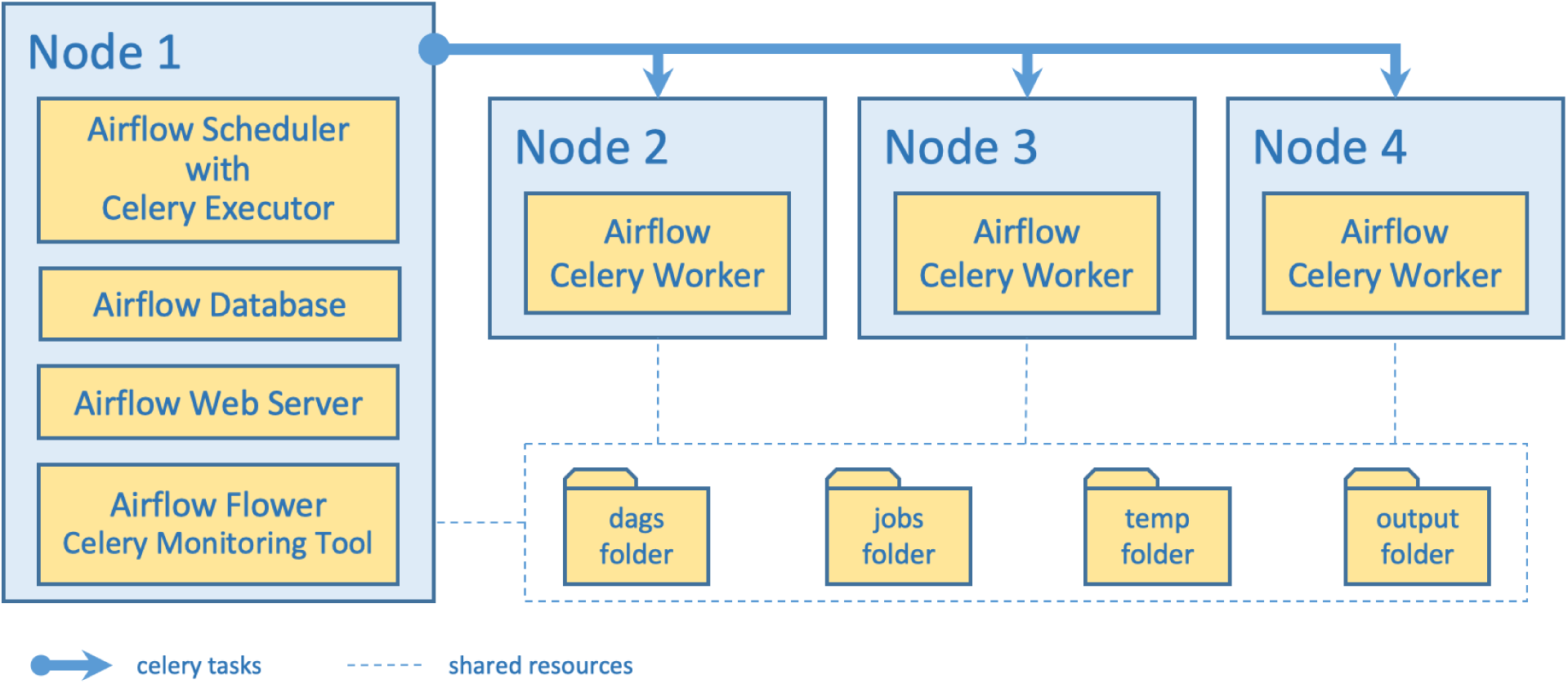
Structure diagram for scaling out CWL-Airflow with Celery cluster of 4 nodes. Node 1 runs the Airflow database to save tasks metadata and the Airflow scheduler with the Celery executor to submit tasks for processing to the Airflow celery workers on nodes 2, 3 and 4. The Airflow and Flower (Celery) web servers allow to monitor and control task execution process. All nodes have shared access to the dags, jobs, temp and output folders.

The example of CWL-Airflow Celery cluster of 4 nodes is shown in Figure 2. The tasks are submitted to the queue by the node 1 and executed by either of the 3 workers (nodes 2, 3 and 4). Node 1 runs two mandatory components - the Airflow database and scheduler. The latter schedules the tasks execution by adding them to the queue. All Celery workers are subscribed to the same task queue. Whenever an arbitrary worker pulls the new task from the queue, it runs it and returns execution results. For the sequential steps, the Airflow scheduler then submits next tasks to the queue. During the task execution intermediate data are kept in the *temp* folder. On successful pipeline completion all output files are moved to the *output* folder. Both *temp* and *output* folders as well as the *dags* and the *jobs* folders are shared between all the nodes of the cluster. Optionally, node 1 can also run the Airflow webserver (Fig. 3) and the Celery monitoring tool Flower (Fig. 4) to provide users with the pipeline execution details.

**Figure 3.**
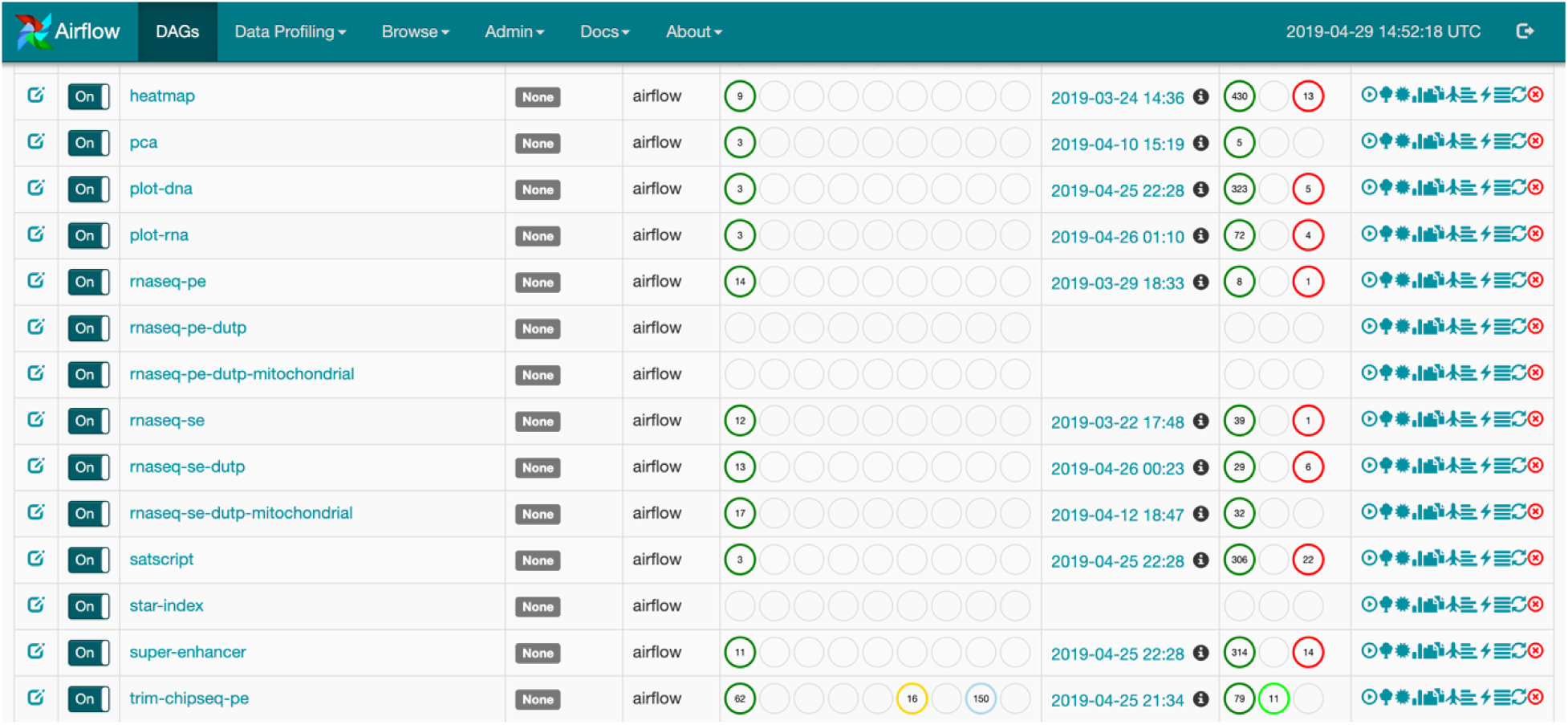
Airflow web interface. The DAGs tab shows the list of the available pipelines, their latest execution dates, number of the active, succeeded and failed runs. The buttons of the right allow to control pipeline execution and obtain additional information on the current workflow and its steps.

**Figure 4.**
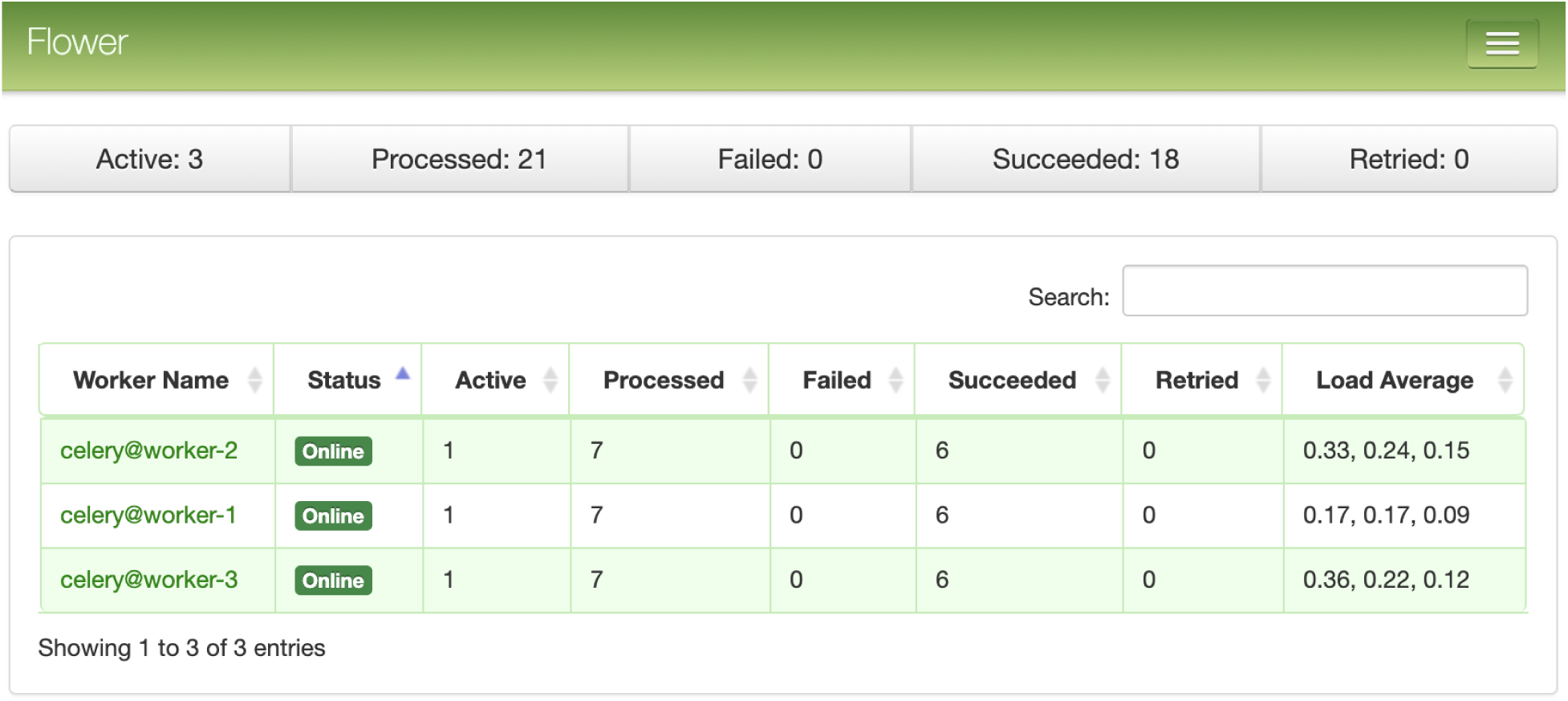
Dashboard of the Celery monitoring tool Flower. Shown are the three Celery workers, their current status and load information.

## Results

### ChIP-Seq analysis with CWL-Airflow

As an example, we used a workflow for basic analysis of ChIP-Seq data [4] (Fig. 5). This workflow is a CWL version of a Python pipeline from BioWardrobe [14,15]. It starts by using BowTie [16] to perform alignment to a reference genome, resulting in an unsorted SAM file. The SAM file is then sorted and indexed with SAMtools [17] to obtain a BAM file and a BAI index. Next MACS2 [18] is used to call peaks and to estimate fragment size. In the last few steps, the coverage by estimated fragments is calculated from the BAM file and is reported in bigWig format (Fig. 5). The pipeline also reports statistics, such as read quality, peak number and base frequency, and other troubleshooting information using tools such as FASTX-Toolkit [19] and BamTools [20]. The directions how to run sample pipeline can be found at [21]. Execution time in CWL-Airflow was similar to that reference implementation (Table 1).

**Figure 5.**
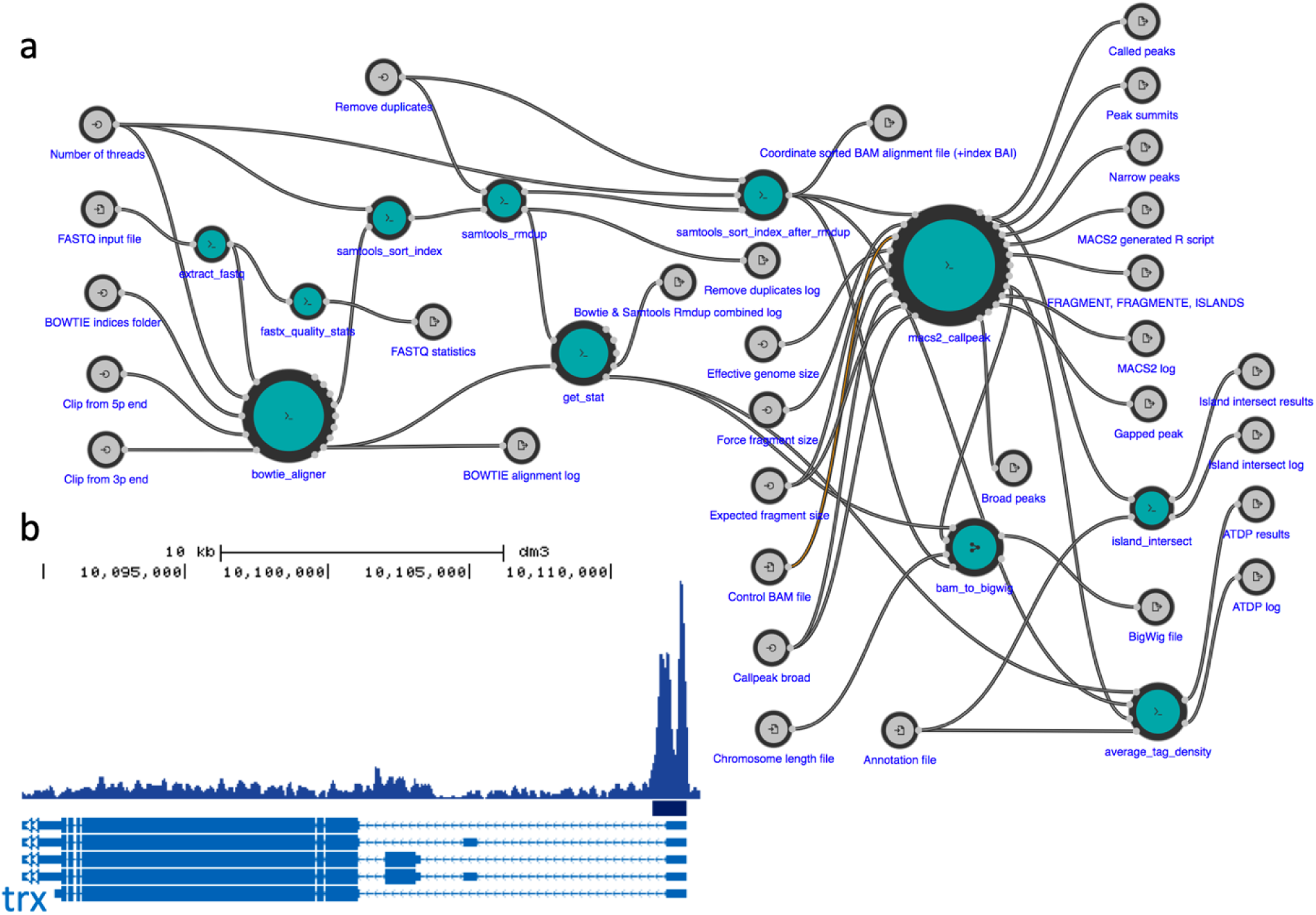
Using CWL-Airflow for analysis of ChIP-Seq data. (a) ChIP-Seq data analysis pipeline visualized by Rabix Composer. (b) Drosophila embryo H3K4me3 ChIP-Seq data (SRR1198790) were processed by our pipeline and CWL-Airflow. UCSC genome browser view of tag density and peaks at trx gene is shown.

The CWL-Airflow package includes two additional demo workflows: (i) identification of super-enhancers [22] and (ii) a simplified version of Xenbase [23] RNA-Seq pipeline. More pipelines can be found elsewhere. In particular, BioWardrobe’s [14] pipelines for analysis of single and paired-end ChIP-Seq, stranded and un-stranded, single and paired RNA-Seq are available on GitHub [24]. Additional collections of tools are available in Rabix Composer [7], a graphical CWL Editor from Seven Bridges and at the dockstore [25].

### Portability of CWL analyses

The key promise of CWL is the portability of analyses. Portability refers to the ability to seamlessly run a containerized CWL pipeline developed for one CWL platform on another CWL platform allowing users to easily share computational workflows. To check whether CWL-Airflow can use pipelines developed by others, we downloaded an alternative workflow for analysis of ChIP-Seq data developed by ENCODE Data Coordination Center [26,27] using a test dataset (CEBPB ChIP-Seq in A549 cells, ENCODE accession: ENCSR000DYI). CWL-Airflow was able to run the pipeline and produced results identical to those obtained with the reference cwl-tool. Execution time is shown in Table 1. As we can see it from the table, running the tested pipelines on the single node CWL-Airflow system resulted in the 18% longer execution time, whereas the 3 node CWL-Airflow cluster reduced execution time by 41% per workflow compared to the reference cwl-tool. These results confirm that CWL-Airflow complies with the CWL specification, support portability and can perform analysis in a reproducible manner. Additional testing of pipeline portability is currently conducted as a part of GA4GH workflow portability challenge [28].

### Using CWL-Airflow in multi-node configuration with Celery executor

To demonstrate the use of CWL-Airflow in a multi-node configuration we set up the Celery cluster of 3 nodes with 4 CPU and 94 GB of RAM each. Every node runs one instance of the Airflow Celery worker. Tasks are queued for execution by the Airflow scheduler that is launched on the first node. Communication between the Celery workers is managed by the message queueing service, RabbitMQ. The latter, as well as the Airflow database and web server, are run on the first node. The results of running two tested pipelines on the Airflow Celery cluster are shown in the Table 1 and show only a slight slow-down on a per-run basis.

## Discussion

CWL-Airflow is one of the first pipeline managers supporting version 1.0 of the CWL standard and provides a robust and user-friendly interface for executing CWL pipelines. Unlike more complicated pipeline managers, the installation of Airflow and the CWL-Airflow extension can be performed with a single *pip install* command. Airflow has multiple advantages compared to the competing pipeline managers (Table 2). Specifically, Airflow provides a wide range of tools for managing workflow execution process, such as pausing and resuming workflow execution, stopping and restarting the individual workflow steps, restarting the workflow from the certain step, skipping part of the workflow by updating the states of the specific steps from a web-based GUI. Similarly, to other workflow management systems, Airflow can run on clusters and the major cloud services. Unlike some of the workflow executors, it supports both Docker and Singularity containerization technologies. The latter is particularly important since many clusters do not allow the use of Docker for security reasons.

**Table 2.** Comparison of the open source workflow managers and engines with existing or planned support for CWL. GUI: graphical user interface; CLI: command line interface; REST API: representational state transfer application programming interface.

| Feature | Airflow & CWL-Airflow |  |  | Rabix |  |  | Toil |  |  | Cromwell |  |  | REANA |  |  | Galaxy |  |  | Arvados |  |  | CWLEXEC |  |  |
| --- | --- | --- | --- | --- | --- | --- | --- | --- | --- | --- | --- | --- | --- | --- | --- | --- | --- | --- | --- | --- | --- | --- | --- | --- |
| Software installation complexity | single python package |  |  | JAR<br>Electron application |  |  | single python package |  |  | JAR |  |  | group of python packages |  |  | group of python packages<br>node.js application |  |  | multiple components for min. 7 nodes system |  |  | JAR |  |  |
| License type | Apache License 2.0 |  |  | Apache License 2.0 |  |  | Apache License 2.0 |  |  | BSD 3-Clause |  |  | MIT License |  |  | AFL v. 3.0 |  |  | Apache License 2.0<br>AGPL-3.0<br>CC-BY-SA-3.0 |  |  | Apache License 2.0 |  |  |
| Workflow description language | CWL v1.0<br>python code |  |  | CWL v1.0 |  |  | CWL v1.0<br>WDL v1.0<br>python code |  |  | CWL v1.0<br>WDL v1.0 |  |  | CWL v1.0<br>Serial<br>Yadage |  |  | XML tool file<br>JSON workflow file |  |  | CWL v1.0 |  |  | CWL v1.0 |  |  |
| Docker containerization | + |  |  | + |  |  | + |  |  | + |  |  | + |  |  | + |  |  | + |  |  | + |  |  |
| Singularity containerization | + |  |  | — |  |  | + |  |  | + |  |  | — |  |  | + |  |  | — |  |  | — |  |  |
| Cloud / cluster processing | + |  |  | — |  |  | + |  |  | + |  |  | + |  |  | + |  |  | + |  |  | + |  |  |
| Workflow execution load balancing<sup>1</sup> | + |  |  | — |  |  | + |  |  | + |  |  | + |  |  | + |  |  | + |  |  | + |  |  |
| Parallel workflow step execution | + |  |  | + |  |  | + |  |  | + |  |  | + |  |  | + |  |  | + |  |  | + |  |  |
|  | GUI | REST API | CLI | GUI | REST API | CLI | GUI | REST API | CLI | GUI | REST API | CLI | GUI | REST API | CLI | GUI | REST API | CLI | GUI | REST API | CLI | GUI | REST API | CLI |
| Add new workflow<sup>2</sup> | — | — | + | + | ∅ | + | ∅ | ∅ | + | ∅ | + | + | ∅ | + | + | + | + | ∅ | + | + | + | ∅ | ∅ | + |
| Set workflow inputs<sup>3</sup> | — | + | + | + | ∅ | + | ∅ | ∅ | + | ∅ | + | + | ∅ | + | + | + | + | ∅ | + | + | + | ∅ | ∅ | + |
| Start / Stop workflow execution | + | + | + | + | ∅ | + | ∅ | ∅ | + | ∅ | + | + | ∅ | + | + | + | + | ∅ | + | + | + | ∅ | ∅ | + |
| Manage workflow execution process<sup>4</sup> | + | + | + | — | ∅ | — | ∅ | ∅ | + | ∅ | — | — | ∅ | + | + | + | + | ∅ | — | + | + | ∅ | ∅ | + |
| Get execution results of the specific workflow<sup>5</sup> | + | — | — | + | ∅ | — | ∅ | ∅ | + | ∅ | + | — | ∅ | + | + | + | + | ∅ | + | + | + | ∅ | ∅ | — |
| View workflow execution logs | + | — | + | + | ∅ | + | ∅ | ∅ | + | ∅ | + | + | ∅ | + | + | + | + | ∅ | + | + | + | ∅ | ∅ | + |
| View workflow execution history and statistics | + | + | + | — | ∅ | — | ∅ | ∅ | + | ∅ | + | — | ∅ | + | + | + | + | ∅ | + | + | + | ∅ | ∅ | + |

<sup>1</sup> assign workflow steps to the different pools and queues; use other resource utilization algorithms provided by the computing environment
<sup>2</sup> load the workflow from the file; create the workflow by combining the steps in GUI
<sup>3</sup> set the path to the job file; set input values through the GUI or CLI
<sup>4</sup> pause/resume workflow execution process; manually restart workflow steps
<sup>5</sup> get output files locations by the workflow id, step id, execution date or other identifiers

Unlike most of the other workflow managers, Airflow provides a convenient web-based GUI that allows to monitor and control pipeline execution. Within this web interface, one can easily track workflow execution history, collect and visualize statistics from multiple workflow runs. Similarly, to some of the other pipeline managers Airflow provides REST API that allows to access its functionality through the dedicated endpoints. The latter can be used by other software to communicate with the Airflow system.

Airflow supports parallel workflow steps execution. Step parallelization can be convenient when the workflow complexity is not that high and the computational resources are not limited. However, when running multiple workflows, especially on a multi-node system, it becomes reasonable to limit parallelism and balance load over the available computing resources. Besides the standard load balancing algorithms provided by the computing environment, Airflow supports pools and queues that allow to evenly distribute tasks among multiple nodes.

Addition of CWL capability to Airflow, has made it more convenient for scientific computing, where the users are more interested in the flow of data rather than the tasks being executed. While Airflow itself (and most of pipeline managers [28]) only define workflows as sequences of steps to be executed (e.g. DAGs), CWL description of inputs and outputs leads to better representation of data flow which allows to better understand data dependencies and produces more readable workflows.

Furthermore, as one of the most lightweight pipeline managers, Airflow contributes only a small amount of overhead to the overall execution of a computational pipeline (Table 1). We believe, however, that this is a small price to pay for the ability to monitor and control workflow execution afforded by Airflow and better reproducibility and portability of biomedical analyses as afforded by the use of CWL. In summary, CWL-Airflow will provide users with the ability to execute CWL workflows anywhere Airflow can run—from a laptop to cluster or cloud environment.

## Abbreviations

CWL: Common Workflow Language
DAG: Directed Acyclic Graph
ChIP-Seq: Chromatin ImmunoPrecipitation - Sequencing
GUI: Graphical User Interface
CLI: Command Line Interface
REST API: Representational State Transfer Application Programming Interface

## Declarations

### Ethics approval and consent to participate

Not applicable.

### Consent to publish

Not applicable.

### Availability of data and materials

No new datasets or materials were generated. The source code is available under Apache license v.2 and can be downloaded from https://barski-lab.github.io/cwl-airflow, http://doi.org/10.5281/zenodo.2669582, RRID: SCR_017196.

### Author contributions statement

AVK and AB conceived the project, AVK and MK wrote the software, MK, AVK and AB wrote and reviewed the manuscript.

## Acknowledgements

The authors thank all members of the CWL working group for their support and Shawna Hottinger for editorial assistance.

## References

1. Leipzig J. A review of bioinformatic pipeline frameworks. Brief Bioinform. 2017;18:530–6.

2. Existing Workflow Systems [Internet]. Available from: https://s.apache.org/existing-workflow-systems

3. Common Workflow Language [Internet]. Available from: http://www.commonwl.org/

4. Docker [Internet]. Available from: https://www.docker.com/why-docker

5. Kurtzer GM, Sochat V, Bauer MW. Singularity: Scientific containers for mobility of compute. PLoS One. 2017;12.

6. Amstutz P, Crusoe MR, Tijanić N, Chapman B, Chilton J, Heuer M, et al. Common Workflow Language, v1.0 [Internet]. Doi.Org. 2016. p. Available from: https://www.commonwl.org/v1.0/Workflow.html

7. Kaushik G, Ivkovic S, Simonovic J, Tijanic N, Davis-Dusenbery B, Kural D. RABIX: an open-source workflow executor supporting recomputability and interoperability of workflow descriptions. Pac Symp Biocomput. 2016;22:154–65.

8. Arvados [Internet]. Available from: https://arvados.org/

9. Giardine B, Riemer C, Hardison RC, Burhans R, Elnitski L, Shah P, et al. Galaxy: a platform for interactive large-scale genome analysis. Genome Res. 2005;15:1451–5.

10. Vivian J, Rao A, Nothaft FA, Ketchum C, Armstrong J, Novak A, et al. Rapid and efficient analysis of 20,000 RNA-seq samples with Toil. bioRxiv. 2016;2:062497.

11. Airflow [Internet]. Available from: http://airflow.incubator.apache.org/

12. Hindman B, Konwinski A, Zaharia M, Ghodsi A, Joseph AD, Katz R, et al. Mesos: A platform for fine-grained resource sharing in the data center. Proc 8th USENIX Conf Networked Syst Des Implement. 2011;295.

13. Celery Project [Internet]. Available from: http://www.celeryproject.org/

14. Kartashov A V, Barski A. BioWardrobe: an integrated platform for analysis of epigenomics and transcriptomics data. Genome Biol. 2015;16:158.

15. Vallabh S, Kartashov A V., Barski A. Analysis of ChIP-Seq and RNA-Seq Data with BioWardrobe. Methods Mol Biol. 2018;1783:343–60.

16. Langmead B, Trapnell C, Pop M, Salzberg SL. Ultrafast and memory-efficient alignment of short DNA sequences to the human genome. Genome Biol. 2009;10:R25.

17. Li H, Handsaker B, Wysoker A, Fennell T, Ruan J, Homer N, et al. The Sequence Alignment/Map format and SAMtools. Bioinformatics. 2009;25:2078–9.

18. Zhang Y, Liu T, Meyer CA, Eeckhoute J, Johnson DS, Bernstein BE, et al. Model-based analysis of ChIP-Seq (MACS). Genome Biol. 2008/09/19. 2008;9:R137.

19. FASTX Toolkit [Internet]. Available from: http://hannonlab.cshl.edu/fastx_toolkit/index.html

20. Barnett DW, Garrison EK, Quinlan AR, Stromberg MP, Marth GT. BamTools: a C++ API and toolkit for analyzing and managing BAM files. Bioinformatics. 2011;27:1691–2.

21. Barski Lab ChIP-Seq SE Workflow [Internet]. Available from: https://barski-lab.github.io/cwl-airflow/#running-sample-chip-seq-se-workflow

22. Hnisz D, Abraham BJ, Lee TI, Lau A, Saint-André V, Sigova A a, et al. Super-enhancers in the control of cell identity and disease. Cell. 2013;155:934–47.

23. Karimi K, Fortriede JD, Lotay VS, Burns KA, Wang DZ, Fisher ME, et al. Xenbase: a genomic, epigenomic and transcriptomic model organism database. Nucleic Acids Res. 2018;46:D861–8.

24. Barski Lab CWL Workflows on GitHub [Internet]. Available from: https://github.com/Barski-lab/workflows

25. O’Connor BD, Yuen D, Chung V, Duncan AG, Liu XK, Patricia J, et al. The Dockstore: enabling modular, community-focused sharing of Docker-based genomics tools and workflows. F1000Research. 2017;6:52.

26. Landt SG, Marinov GK, Kundaje A, Kheradpour P, Pauli F, Batzoglou S, et al. ChIP-seq guidelines and practices of the ENCODE and modENCODE consortia. Genome Res. 2012;22:1813–31.

27. ENCODE ChIP-Seq pipeline [Internet]. Available from: https://github.com/ENCODE-DCC/pipeline-container

28. GA4GH-DREAM Workflow Execution Challenge [Internet]. Available from: https://www.synapse.org/#!Synapse:syn8507133/wiki/415976

